# Refinement of highly flexible protein structures using simulation-guided spectroscopy

**DOI:** 10.1101/319335

**Authors:** Jennifer M. Hays, Marissa K. Kieber, Jason Z. Li, Ji In Han, Linda Columbus, Peter M. Kasson

## Abstract

Highly flexible proteins present a special challenge for structure determination because they are multi-structured yet not disordered, and the resulting conformational ensembles are essential for understanding function. Determining such ensembles is difficult because many measurements that capture multiple conformational populations provide sparse data. A powerful opportunity exists to leverage molecular simulations for spectroscopic experiment selection. We have developed an information-theoretic approach to guide experiments by identifying which measurements best refine the underlying conformational ensemble. We have tested this approach on three flexible bacterial proteins. For proteins where a clear mechanistic hypothesis drives label selection, our approach systematically identifies labels that would test this hypothesis. Furthermore, when available data do not yield an obvious mechanistically-guided label selection strategy, our approach guides label selection and produces conformational refinement that significantly outperforms standard structure-guided approaches. Our information-theoretic approach to label selection thus offers a particular advantage when refining challenging, underdetermined protein conformational ensembles.

---

Heterogeneous conformational ensembles play critical roles in molecular recognition and cellular regulation, yet high-resolution structure determination has typically required reducing these ensembles to only a few states. Since the full equilibrium ensemble is often key to understanding biochemical function, alternative experimental techniques have been developed to probe the full ensemble distribution rather than either a few low-energy states or an equilibrium average^[1]^. However, these experiments measure only a small number of atomic degrees of freedom^[2]^: for instance, double electron-electron resonance (DEER) and single-molecule Förster Resonance Energy Transfer (smFRET) spectroscopy, which utilize pairs of labeled amino acids, typically provide data for ~10 measurements per system. Thus, experiment selection is currently the limiting factor in how much information can be obtained on an ensemble.

Prior quantitative approaches to experiment selection have relied on pre-existing high-resolution structural and kinetic models. Two recent studies have shown, retrospectively, that leveraging either Markov State Models^[3]^ or normal modes calculated from elastic network models^[4]^ can select good labels for DEER experiments. But for systems where traditional structural or kinetic models are incomplete or fundamentally underdetermined due to conformational flexibility, it remains extremely challenging to determine which pairs of residues should be chosen for labeling. Indeed, for highly flexible systems, experiment selection is often guided by the limitations of the experimental modality rather than the system under study. We have therefore developed a general, information-theoretic formalism for selecting optimal spectroscopic experiments. We briefly summarize the theory below, then discuss the application of this method to three conformationally heterogeneous bacterial proteins.

An optimal set of spectroscopic experiments has two properties: each experiment yields the maximum amount of information on the conformational ensemble and minimally redundant information with other experiments in the set to avoid wasting labeling and measurement effort (Fig 1). The maximum-relevance, minimum redundancy (mRMR) algorithm exactly satisfies these criteria^[5a, 5b.]^ To select a set of *N* spectroscopic experiments, we maximize the mutual information (MI) between the set of spectroscopic observables {*O_i_*} and the conformation *C*:

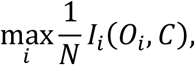

simultaneously minimizing the pairwise MI between spectroscopic variables *O_i_* and *O_j_* (Fig 1):

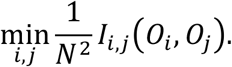

where *I*(*X*,*Y*) is the mutual information between random variables *X* and *Y*:

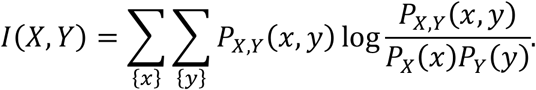

**Figure 1.**
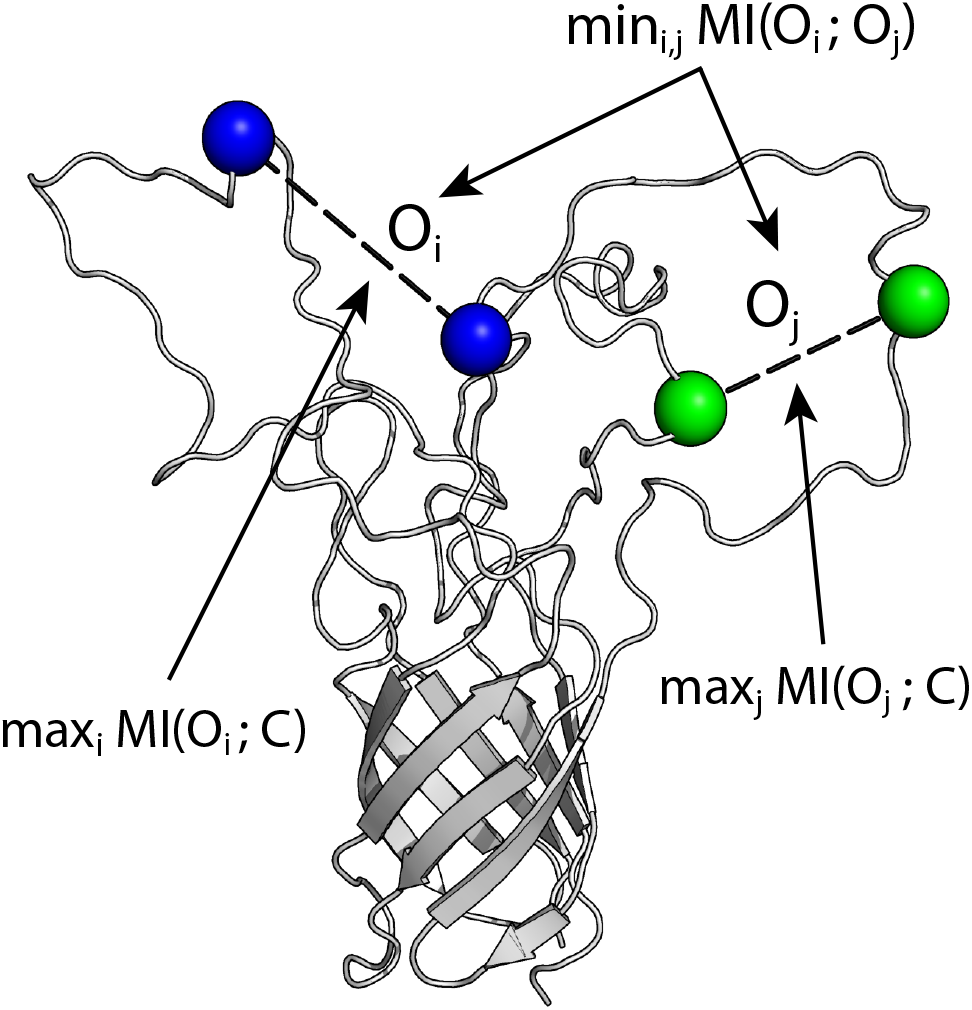
The maximum-relevancy, minimum-redundancy (mRMR) method applied to spectroscopic observables {*O_i_*}. The optimal set of spectroscopic experiments that report on variables {*O_i_*} are maximally informative of the conformation *C* and minimally redundant with each other. Informativeness and redundancy are quantified via mutual information (MI).

This method is particularly useful for flexible proteins that resist traditional structure determination because it identifies, *by design*, those observables which are maximally underdetermined.

In our study, we make two approximations: we use deliberately undersampled estimates of the protein conformational ensemble to select labels for further refinement, and we approximate the spectroscopic variable O_i_ as the C_α_-C_α_ pairwise distance distribution between labeled residues. The first approximation is critical to the success of our method: by identifying underdetermined degrees of freedom we can improve an incomplete estimate of the conformational ensemble rather than requiring a well-sampled starting model. The second approximation is an implementation rather than theoretical concern and we will discuss how it can be removed. The success of the mRMR method and these approximations is demonstrated below on a set of flexible bacterial outer membrane proteins.

Beta-barrel membrane proteins are excellent candidates for the mRMR approach because many contain flexible regions that are difficult to characterize experimentally yet have regions of secondary structure that make spectroscopic experiments tractable^[6]^. We have performed molecular dynamics (MD) simulations on three bacterial outer membrane proteins and applied the mRMR algorithm to select optimal DEER experiments. We have chosen FhuA, an *E. coli* iron transporter^[7]^, OprG, a Pseudomonal small-molecule transporter^[8]^, and Opa_60_, a Neisserial virulence-associated protein that binds cell-surface proteins but does not function as a transporter^[9]^. The FhuA conformational ensemble has been characterized via DEER experiments guided by pre-existing mechanistic hypotheses^[10]^; it is thus a good test system for determining whether the mRMR algorithm identifies similar labels to those identified by spectroscopists. OprG, a more challenging system, has been studied using a combination of NMR and mutational experiments,^[6c]^ but the mechanisms by which transport is regulated remain unknown. Finally, Opa_60_ represents a particularly challenging system since it displays substantial, experimentally underdetermined conformational flexibility that controls its binding mechanism^[6b]^. We have therefore studied this final system prospectively: choosing a set of residue-residue pairs using the mRMR algorithm, measuring them with DEER, incorporating the experimental data into MD simulation, and evaluating this ensemble versus one refined with spectroscopist-selected pairs (SSP).

For each protein, we generated initial estimates of the conformational ensembles using ensemble MD simulations that were deliberately undersampled at 2 *μ*s per protein. We used the mRMR algorithm on these data to select sets of pairwise distances that optimally report on undersampled regions of phase space (Fig 2).

**Figure 2.**
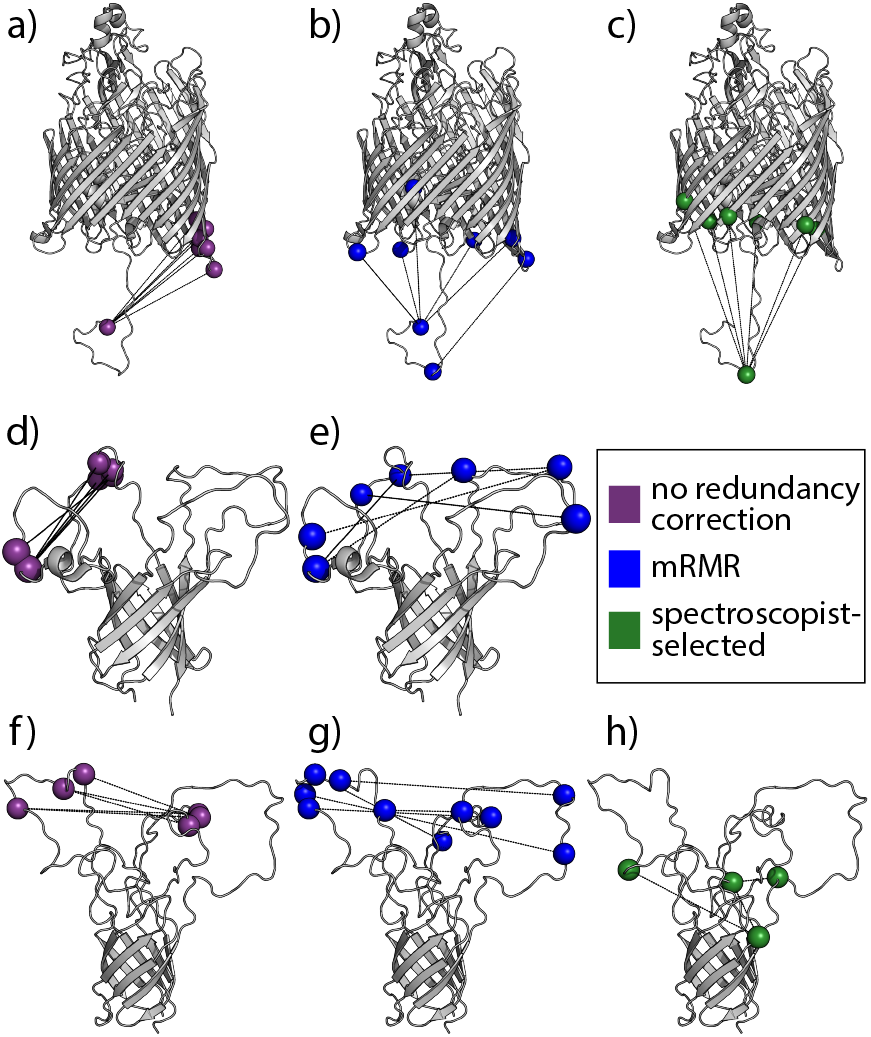
mRMR captures highly informative residues that are minimally redundant for three bacterial outer membrane proteins. Selection via mutual information alone yields informative but redundant pairs (magenta). Selection via mRMR (blue) removes this redundancy. These residues are better distributed across the structures of all three bacterial proteins than the top-ranking pairs via MI alone or pairs selected by spectroscopists (green) according to current practice in the field. Panels a-c, d-e, and f-h show residues selected for FhuA, OprG, and Opa, respectively. Spectroscopists had previously selected and performed DEER measurements on FhuA and Opa.

In the case of FhuA, spectroscopists selected label pairs near the N-terminal domain interaction motif, which is conformationally heterogeneous and regulates transport, and the periplasmic side of the beta-barrel, using a standard triangulation strategy (Fig. 2c)^[10]^. It is encouraging that mRMR-based selection identifies similar residues (Fig. 2a), with the addition of one pair spanning just the N-terminal domain. Predicted label pairs not only spanned these distances between the N-terminal domain and the barrel, but they also specifically identified distances between this domain and one side of the barrel as most informative. Analysis of the DEER data independently identified this side of the barrel as interacting with the N-terminal domain. These two findings on FhuA, a relatively well-understood transport protein, show that the mRMR method can select label pairs that reflect best spectroscopic understanding and yield insight into conformational heterogeneity.

Our method provides even greater potential benefit when less is known about transport mechanism, as in the case of OprG, and may be useful in verifying claims of loop involvement in OprG transport. Both the mechanism and the substrates for OprG transport are unclear: OprG may transport small, hydrophobic compounds via a lateral gating mechanism or small amino acids via the barrel channel; OprG crystal structures support the former hypothesis^[11]^, while recent NMR and mutational studies suggest the latter^[6c]^. Non-transporting mutants studied with NMR generally have more ordered loops. One loop containing a short helical region had especially restricted motion, suggesting that the dynamics of this loop are critical to transport. Interestingly, this loop participates in all five residue-residue pairs that are most informative on the OprG conformation (Fig 2d) and in three of the five top-scoring mRMR pairs (Fig 2e). Thus, mRMR analysis yields label pairs that lend support to existing mechanistic hypotheses and, most importantly, identify experiments to test these hypotheses.

As a robust test of mRMR-based label selection, we prospectively tested its ability to select DEER experiments to refine the conformational ensemble of Opa_60_, the most challenging protein in our evaluation set. DEER data were acquired using label pairs selected via both mRMR and traditional structure-based selection, and we assessed the relative utility of each method in refining the ensemble. Opa_60_’s long, flexible loops are both critical for function^[6a, 6b]^ and challenging for previous DEER pair selection methods. In contrast to FhuA or OprG, no structural or functional data provide strong guidance on which loop residues are responsible for function, in this case receptor engagement. Prior hybrid NMR-MD refinement of the *apo* conformational ensemble did not provide sufficient insight into the binding mechanism. Normal-mode approaches developed by Zheng and Brooks have been applied to identify informative, non-redundant label sets for DEER that differentiate pairs of structures when such structural data do exist^[4, 12]^, but this is not the case for Opa_60_. Furthermore, normal-mode calculations from an Opa_60_ elastic network model do not correlate with loop flexibility measured via NMR relaxation timescales (Fig S1). Thus, spectroscopists are left with more than 5,000 possible inter-loop pairs to choose from. In the following discussion, we demonstrate that mRMR selection method radically improves structural refinement compared to standard spectroscopic practice for systems that were previously intractable.

We prospectively tested mRMR pair selection by refining the Opa_60_ conformational ensemble using two independently identified label sets: one selected using the mRMR algorithm and the other independently chosen by spectroscopists. The top five top-scoring mRMR pairs span multiple combinations of inter-loop distances, and the top ten pairs capture all possible combinations of the loops (Fig. 2g). By contrast, the top ten pairs identified using maximum relevancy alone span a single loop-loop pair, likely losing important information about the dynamics of the other loop (Fig. 2f). The spectroscopist-selected pairs are primarily short barrel-loop distances, largely because the length of the loops permits distances too long to be measured via DEER, so spectroscopic best practice is to select a more conservative set of pairs. However, even had a spectroscopist decided to measure loop-loop distances, the chance of selecting a pair within the top 25% of pairs identified via mRMR would be only 7%, showing a strong advantage for the systematic selection methods developed here.

Because Opa_60_ is so conformationally flexible, approximating the label-label distance distributions as C_α_-C_α_ distributions introduces negligible error relative to the backbone motions of the protein. However, label flexibility becomes increasingly important to label selection as protein flexibility decreases. If label conformations are an important factor in site-selection, explicit labels may be introduced as follows. First, unrestrained simulations of the wild-type protein are used to calculate initial mRMR estimates. Explicit labels are introduced for each top-ranked residue-residue pair, and one additional simulation is performed per pair. The mRMR scores are recalculated for each simulation to determine the effects of label side-chain conformation on the final mRMR rankings. A “forward model” can also be introduced for the spectroscopic observable, such as the predicted DEER spectrum^[12b]^, using the explicit-label simulations.

To assess the quality of mRMR-guided versus structure-guided refinement, we estimated the Opa_60_ conformational ensemble using DEER data on pairs selected via each approach (see Methods). We then compared the resulting ensembles using two independent metrics which we developed to quantitatively evaluate “quality of refinement.” As a first metric, we measured the ability of each refined ensemble to predict DEER data that were held back from refinement as a test set. Refinement using mRMR-selected label pairs yielded significantly better agreement with the test DEER data: seven of the eight test distributions are better captured by the mRMR-guided ensemble than the structure-guided ensemble (Fig 3).

**Figure 3.**
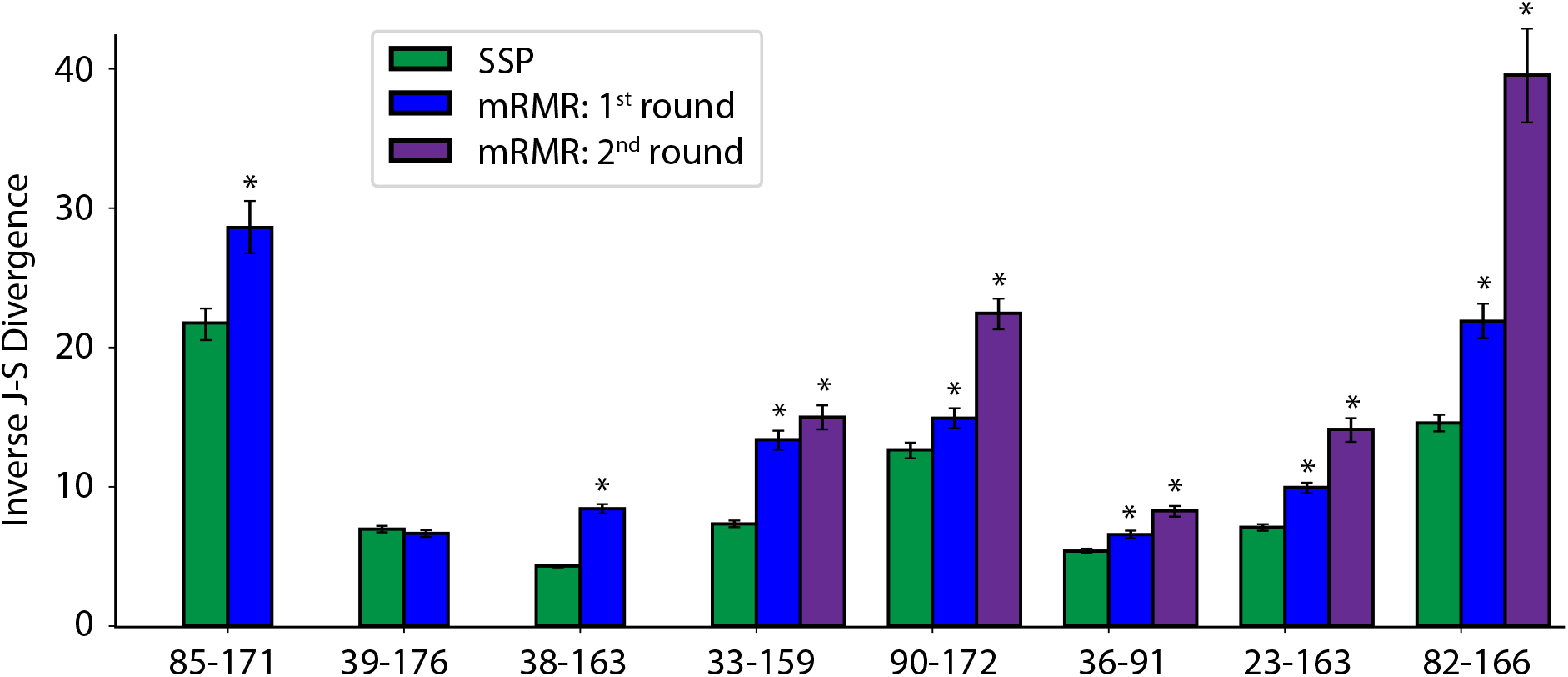
mRMR-guided refinement better predicts test DEER distributions than structure-guided refinement. Each refined Opa_60_ ensemble was evaluated by its ability to predict 8 residue-residue pairs identified via mRMR after the first round of refinement and measured using DEER. Conformational ensembles refined using mRMR-selected pairs predict these DEER distributions significantly better than conformational ensembles refined using spectroscopist-selected pairs (SSP) in seven of eight cases, quantified as inverse Jensen-Shannon divergences. Three of these DEER pairs were used for a second round of mRMR refinement; the resulting conformational ensemble out-performs both 1^st^-round ensembles in predicting the five pairs not used for refinement. Error bars represent 90% confidence using 1000 bootstrap replicates, and * denotes p < 0.01 via two-tailed t-tests.

We also analyzed the dimensionality of the conformational ensembles obtained from refinement using structured-guided versus mRMR-guided DEER data. Given sufficient sampling, a better-refined conformational ensemble will have lower dimensionality, approaching the “true” ensemble in the lower limit. We therefore developed a quantitative measure for the dimensionality of a conformational ensemble (see Methods). Because residue-residue distances yield an overcomplete basis set, we lumped together highly related distance variables at different thresholds of relatedness (ε) and calculated the number of independent distance variables required to describe the ensemble at each threshold. At every threshold tested, refinement with mRMR-selected DEER data yielded a conformational ensemble of lower dimensionality than refinement with spectroscopist-selected DEER data (Fig. 4). This indicates that DEER data from mRMR-selected pairs refine the conformational ensemble more efficiently than data from pairs selected according to current state of the art.

**Figure 4.**
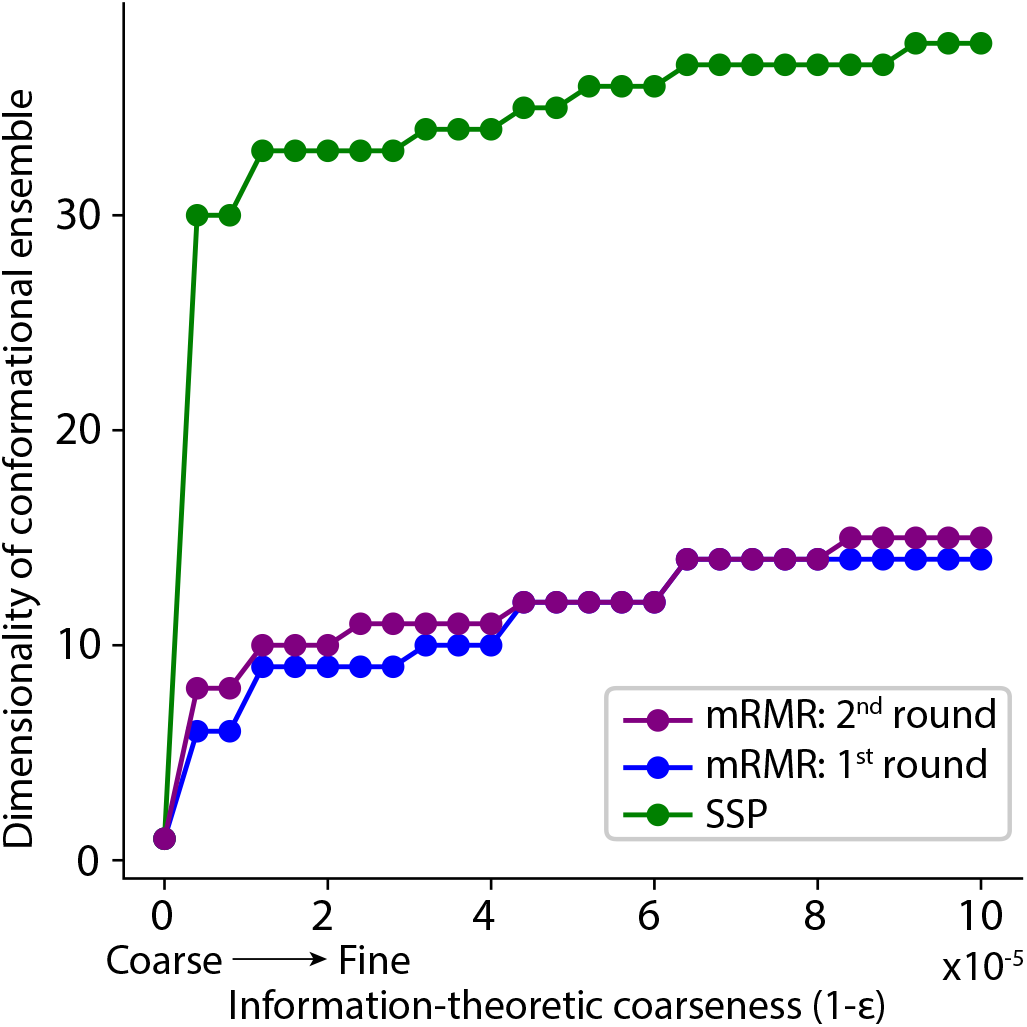
mRMR-guided refinement produces ensembles of lower dimensionality than structure-guided refinement. To assess convergence of the conformational ensemble, we calculated the dimensionality of the conformational ensemble at each information theoretic resolution ε (details in Supplement). Ensembles refined using mRMR-selected pairs are of lower dimensionality than ensembles refined using spectroscopist-selected pairs (SSP) by 20-25.

These tests demonstrate that mRMR provides a robust approach to spectroscopic label selection, particularly for flexible proteins where structural estimates are more challenging and the difference in data quality between optimally selected labels and poorly selected labels is greater. When strong mechanistic hypotheses have guided prior DEER experiments, mRMR yields label pairs that would test these hypotheses. For proteins such as Opa_60_ where mechanistic understanding is insufficient to guide experiment selection, we show via prospective testing that mRMR selection outperforms unaided spectroscopists. Therefore, we believe that mRMR will be of general use in guiding spectroscopic experiment selection for DEER and for other label-based spectroscopy methods such as single-molecule FRET and paramagnetic resonance enhancement. The method can be further extended to differentiate mechanistic hypotheses rather than conformations. For systems like OprG where two mechanistic hypotheses exist, mRMR could be used to identify which spectroscopic variables optimally distinguish conformational features *specific* to one mechanism or the other. Conformational flexibility and heterogeneity are some of the most challenging and exciting frontiers in understanding protein structures, and mRMR will increase the ability of these experimental methods to efficiently refine such conformational ensembles.

## Supporting Information

The Supporting Information is available free of charge on the ACS Publications website at [doi]. Detailed computational and experimental methods are provided as well as validation measures. (PDF)

## AUTHOR INFORMATION

The authors declare no competing financial interests.

## ACKNOWLEDGMENT

The authors thank G. Cortina, D. Cafiso, and L. Tamm for many helpful discussions. Computational resources were provided by XSEDE Comet at San Diego Supercomputing Center through allocation MCB150128. Additional resources were provided by NCSA Blue Waters, SNIC and the Center for Parallel Computation at KTH. This work was supported by NIGMS grants R01GM087828 to L.C. and R01GM115790 to P.M.K., a Wallenberg Academy Fellowship to P.M.K., a Blue Waters Fellowship (OCI-0725070) to J.M.H., and by a collaborative award from the Jeffress Trust.

## REFERENCES

[1] aP. Bernado, E. Mylonas, M. V. Petoukhov, M. Blackledge, D. I. Svergun, J Am Chem Soc 2007, 129, 5656-5664; bG. Wei, W. Xi, R. Nussinov, B. Ma, Chem Rev 2016, 116, 6516-6551; cE. J. Levin, D. A. Kondrashov, G. E. Wesenberg, G. N. Phillips, Jr., Structure 2007, 15, 1040-1052; dA. M. Bonvin, A. T. Brunger, J Mol Biol 1995, 250, 80-93.

[2] aG. Jeschke, Proteins 2016, 84, 544-560; bR. Ward, M. Zoltner, L. Beer, H. El Mkami, I. R. Henderson, T. Palmer, D. G. Norman, Structure 2009, 17, 1187-1194; cJ. N. Rao, C. C. Jao, B. G. Hegde, R. Langen, T. S. Ulmer, J Am Chem Soc 2010, 132, 8657-8668; dS. J. Hirst, N. Alexander, H. S. McHaourab, J. Meiler, J Struct Biol 2011, 173, 506-514.

[3] S. Mittal, D. Shukla, J Phys Chem B 2017.

[4] G. Jeschke, J Chem Theory Comput 2012, 8, 3854-3863.

[5] aH. Peng, F. Long, C. Ding, IEEE Transactions on pattern analysis and machine intelligence 2005, 27, 1226-1238; bC. Ding, H. Peng, J Bioinform Comput Biol 2005, 3, 185-205.

[6] aJ. N. Martin, L. M. Ball, T. L. Solomon, A. H. Dewald, A. K. Criss, L. Columbus, Biochemistry 2016, 55, 4286-4294; bD. A. Fox, P. Larsson, R. H. Lo, B. M. Kroncke, P. M. Kasson, L. Columbus, J Am Chem Soc 2014, 136, 9938-9946; cI. Kucharska, P. Seelheim, T. Edrington, B. Liang, L. K. Tamm, Structure 2015, 23, 2234-2245; dA. Pautsch, G. E. Schulz, Nat Struct Biol 1998, 5, 1013-1017; eA. Arora, F. Abildgaard, J. H. Bushweller, L. K. Tamm, Nat Struct Biol 2001, 8, 334-338.

[7] G. S. Moeck, J. W. Coulton, K. Postle, J Biol Chem 1997, 272, 28391-28397.

[8] R. E. Hancock, F. S. Brinkman, Annu Rev Microbiol 2002, 56, 17-38.

[9] S. E. McCaw, E. H. Liao, S. D. Gray-Owen, Infect Immun 2004, 72, 2742-2752.

[10] J. L. Sarver, M. Zhang, L. Liu, D. Nyenhuis, D. S. Cafiso, Biochemistry 2018, 57, 1045-1053.

[11] D. S. Touw, D. R. Patel, B. van den Berg, PLoS One 2010, 5, e15016.

[12] aW. Zheng, B. R. Brooks, Biophys J 2005, 88, 3109-3117; bG. Jeschke, Protein Sci 2017.

